# Macrophages undergo necroptosis during severe influenza A infection and contribute to virus-associated cytokine storm

**DOI:** 10.1101/2022.04.20.488871

**Authors:** André C. Ferreira, Carolina Q. Sacramento, Filipe S. Pereira-Dutra, Natália Fintelman-Rodrigues, Priscila P. Silva, Mayara Mattos, Caroline S. de Freitas, Andressa Marttorelli, Gabrielle R. de Melo, Mariana C. Macedo, Isaclaudia G. Azevedo-Quintanilha, Aluana S. Carlos, João Vitor Emídio, Cristiana C. Garcia, Patrícia T. Bozza, Fernando A. Bozza, Thiago M. L. Souza

**Author notes:** The authors contributed equally as the first authors. The authors contributed equally as the last authors.

## Abstract

Influenza A virus (IAV) causes a major public health concern, because it is one of the leading causes of respiratory tract infections and hospitalization. Severe influenza has been associated with the cytokine storm, and IAV productive infection cell death in airway epithelial cells may contribute to the exacerbation of this proinflammatory event. On the other hand, IAV replication in macrophages is non-permissive and whether this immune cell may contribute to severe influenza physiopathology requires more details. Here, we investigated IAV-induced macrophage death, along with potential therapeutic intervention. We found that IAV or simply its surface glycoprotein hemagglutinin (HA) triggers necroptosis in human and murine macrophages in a Toll-like receptor-4 (TLR4) and TNF-dependent manner. Anti-TNF treatment, with the clinically approved drug etanercept, prevented the engagement of the necroptotic loop and mice mortality. impaired IAV-induced pro-inflammatory cytokine storm and lung injury. Our results implicate macrophage necroptosis with severe influenza in experimental models and potentially repurpose a clinically available therapy.

**Author Summary:** Various fates of cell death have an integral role in the influenza A virus (IAV) pathogenesis and lung/respiratory dysfunction. IAV physiopathology is not restricted to airway epithelial cells, where this virus actively replicated. Macrophages should support both viral clearance and priming of adaptative immune response in patients that adequately control influenza. However, during severe IAV infection, macrophages – which are unable to support a permissive viral replication - undergo disruptive cell death and contribute to the exacerbated production of proinflammatory cytokines/chemokines. We characterized this process by showing that IAV or just its surface glycoprotein hemagglutinin (HA) trigger necroptosis, a disruptive and TNF-dependent cells death. Since TNF is a hallmark of pro-inflammatory cell death, we blocked this mediator with a repurposed biomedicine etanercept, which prevented the severe IAV infection in the experimental model. The present work improves the knowledge of influenza pathophysiology by highlighting the importance of macrophage cell death during severe infection.

## Introduction

Acute respiratory illness caused by influenza A (IAV) is associated with epidemic [1] and pandemic outbreaks, like during the emergence of Influenza A(H1N1)pdm09during 2009 [2]. Besides higher incidence of complicated clinical outcomes during pandemic situations, seasonal influenza also represents a public health threat. Influenza presentation ranges from mild to potentially lethal cases of severe acute respiratory syndrome (SARS) [3,4]. Annually, there are an estimated 1 billion cases of influenza, of which 3–5 million are severe and 10 to 20 % of these complicated cases may result in deaths [4–6].

IAV has a negative-sense and segmented RNA genome coding for at least 11 viral proteins, of which hemagglutinin (HA) and neuraminidase (NA) are the major surface glycoproteins [7]. During its life cycle, HA attaches to sialic acid residues on cellular surface, allowing IAV to be endocytosed [8]. After, HA-dependent fusion of viral envelope with endosome membrane, viral RNA and associated proteins are transported to the cellular nucleus, where replication and transcription take place. Newly synthesized proteins are assembled, virions bud the cell membrane and NA assists their final release [9]. During this process, infected cells may succumb to death, and the pathway to death has been associated to IAV pathogenesis [9,10]. In airway epithelial cells, where IAV performs its replicative cycle completely and generates infectious progeny, infection activates the receptor interaction serine/threonine-protein kinases 1 and 3 (RIPK1 and RIPK3) promoting apoptosis and necroptosis [11–14]. Whereas apoptosis has been proposed as cell death related to limit IAV-induced tissue damage [15], virus-induced necrotic cell death may exacerbate inflammatory engagement [14,16]. During necroptosis, cells are triggered to shift from an apoptotic-like cell death to the disruption of the plasma membrane, facilitating the release of immunomodulatory damage associated with molecular patterns (DAMPs), leading to inflammation [17]. The pro-inflammatory cytokine storm is an important hallmark of IAV-induced severe pneumonia, strongly associated with lung/respiratory dysfunction [18].

Moreover, IAV physiopathology is not restricted to airway epithelial cells. Macrophages are also exposed to this virus, although they are unable to harbor a permissive virus replication[19]. In the IAV-infected lung, both resident macrophages by the infected and macrophages derived from blood monocytes, since they migrate towards the respiratory tract and differentiate to macrophages [20,21]. To limit the infection in the lower respiratory tract, macrophages should clear the virus particles and prime the adaptive immune response [22]. However, IAV-infected macrophage death has been related to overproduction of local and systemic cytokines/chemokines, cellular infiltration, extracellular matrix degradation, and airway epithelial denudation. Whether these events are influenced by the mechanism of macrophage death require more details [22]. Here, we show that IAV- and HA-induced necroptosis in primary macrophages lead to a loop of events involving Toll-like receptor 4 (TLR4), Tumor necrosis factor (TNF), and RIPK1, resulting in exacerbation of inflammatory response. Moreover, we show that the blocked TNF-related necroptosis signaling by biodrug etanercept increased the survival of IAV-infected animals and jeopardized the inflammatory response associated with several IAV infections.

## RESULTS

### IAV and its HA induce necroptosis and pro-inflammatory response in primary macrophages

For initial assessments of IAV-induced macrophage death, we evaluated the cell surface exposure of phosphatidylserine (PS) and the loss of plasma membrane integrity, by respectively labeling the cells with AnnexinV and propidium iodide (PI) through flow cytometry. We observed that IAV increased the number of AnnexinV^+^ and AnnexinV^+^/PI^+^ cells **(Figure 1A)**. Because IAV life cycle in macrophages is generally dead-end [19], we tested whether the viral protein responsible for initial attachment and a known pathogen-associated molecular patterns (PAMPs)[23], the HA, could also trigger a similar cell fate. Indeed, HA also enhanced the number of AnnexinV^+^ and AnnexinV^+^/PI^+^ macrophages **(Figure 1A)**. Next, we treated the murine macrophages cells with pharmacological inhibitors of necroptosis (Nec-1, an inhibitor of RIPK1) or apoptosis (zVAD, a pan-caspase inhibitor) before IAV infection. Nec-1 pretreatment prevented the increase of annexinV^+^/PI^+^ cellular content **(Figure 1B and Figure S3)** and LDH levels quantified in the culture supernatant **(Figure 1C),** whereas zVAD was not able to prevent these events. The same results were obtained with human macrophages, because Nec-1 prevented IAV- and HA-induced enhancement of LDH levels **(Figure 1D)**. Of note, TNF was used as a positive control to induce apoptosis [24], and TNF/zVAD as a positive control of necroptosis [25]. Consistently with cytometry findings and LDH levels, IAV and HA significantly increased the expression of p-RIPK1/RIPK3 that transduce the necroptotic signal to the effector protein Mixed lineage kinase domain-like (MLKL) **(Figure 1E and Figure S4)**, the pore-forming protein involved with membrane disruption [10,26]. Importantly, pretreatment with Nec-1 prevented IAV- and HA-induced engagement of necroptotic signals in macrophages **(Figure 1E and Figure S4)**.

**Figure 1.**
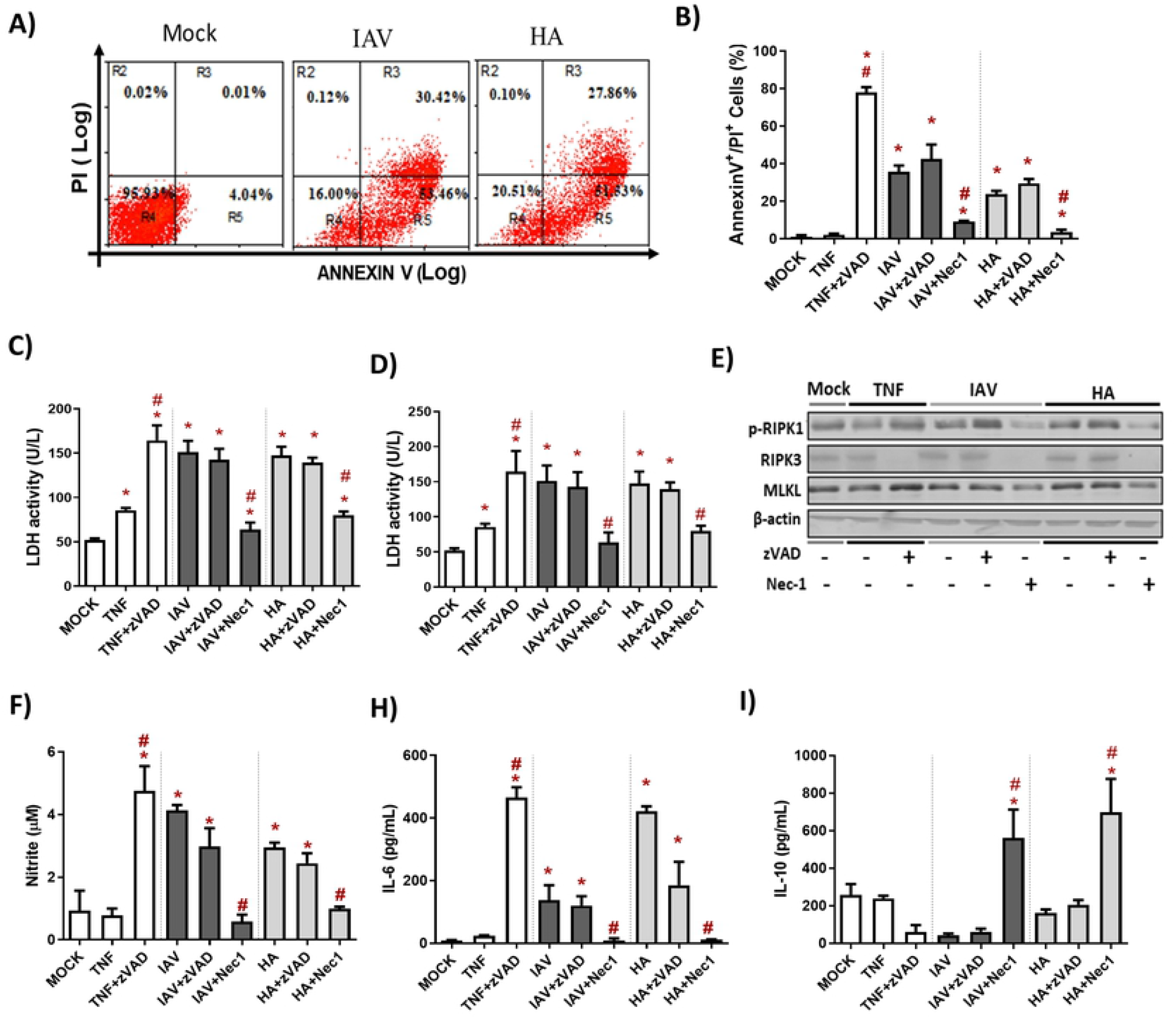
IAV and viral HA induce necroptosis and pro-inflammatory response in primary macrophages. Murine (BMDM) or human macrophages (MDM) cultures were pretreated or not with zVAD (10 μM) or Nec −1 (25 μM) and infected with IAV at MOI of 0.25, exposed to the viral 10 ng/mL of HA or 1 ng/mL of TNF-α. (A-B) BMDM cells death was also evaluated by flow cytometry analysis of AnnexinV/PI positive cells. Assessment of cell viability through the measurement of LDH release in the supernatant of BMDM (C) or human (D). (E) The expression of p-RIPK1, RIPK3 and MLKL were detected in BMDM lysates by Western blotting. β-actin levels were used for control of protein loading. (F) Levels of nitrite were measured by Griess method in supernatant of BMDM cultures after 24h of stimulus. The levels of (H) IL-6 and (K) IL-10 were measured by ELISA assay in supernatant of BMDM cultures after 24h of stimulus. Data are presented as the mean ± SEM of 5 independent experiments * P < 0.05 versus control group (MOCK); # P < 0.05 versus respective untreated infected/stimulated group.

Because necroptosis is associated with the pro-inflammatory response, macrophages infected with IAV or exposed to HA produced nitric oxide **(Figure 1F)** and IL-6 **(Figure 1G)**. Nec-1 pretreatment on these cells completely shifted the macrophage phenotype to prevent the pro-inflammatory phenotype and increasing the content of the regulatory cytokine IL-10 (**Figure 1F-I**). Together, these data suggest that IAV-induced necroptosis in macrophages may be directly related to the pro-inflammatory cytokine storm observed in severe disease.

### Toll-like receptor 4 (TLR4) is engaged during IAV- and HA-induced necroptosis

Because influenza HA has been shown to engage TLR4 as an early signal of innate immune response in leukocytes [27,28], we hypothesized that this event could also occur in macrophages as an upstream event to trigger necroptosis. Thus, macrophages were pre-treated with an TLR4 signaling pathway inhibitor (CLI95) and exposed to IAV or HA. CLI95-treated IAV/HA-exposed macrophages display lower levels of LDH **(Figure 2A)** and annexinV^+^/PI^+^ cells **(Figure 2B)**, compared to untreated control macrophages **(ctr; Figure 2)**.

**Figure 2.**
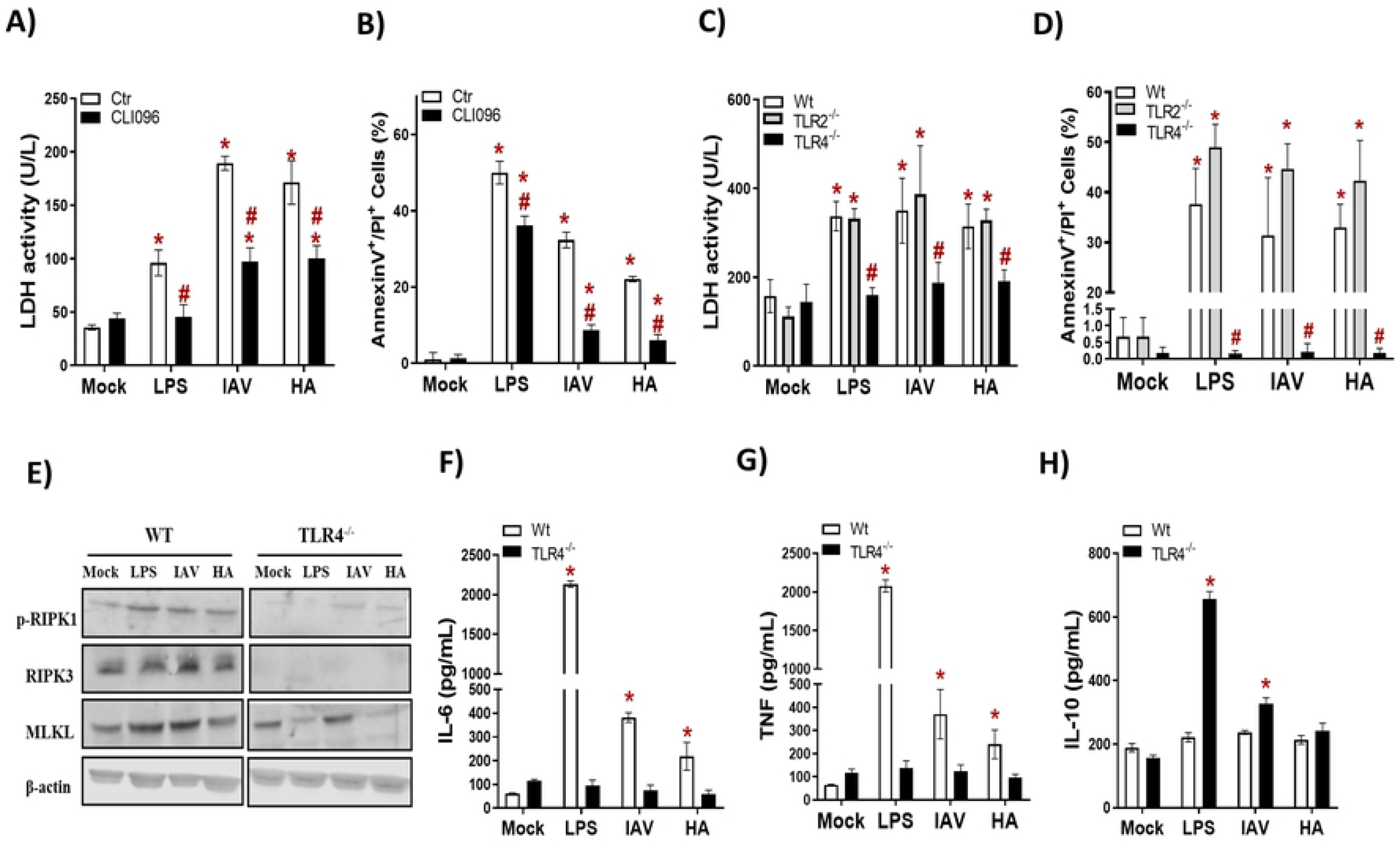
TLR4 engagement is necessary to IAV- and HA-induced necroptosis. **(A-B)** BMDM cultures were pretreated with 2 μM of CLI95 for 2 h and infected with IAV at MOI of 0.25, exposed to 10 ng/mL of HA or LPS. After 24 h, LDH was measured (A), annexinV/PI labeled (B). * P < 0.05 versus respective control group (MOCK); # P < 0.05 versus respective untreated infected/stimulated group. (C-D) BMDM cultures from WT, TLR2^-/-^ and TLR4^-/-^ mices were infected with IAV at MOI of 0.25, exposed to 10 ng/mL of HA or LPS. After 24 h, LDH was measured (C), annexinV/PI labeled (D). (E) The expression of p-RIPK1, RIPK3 and MLKL were detected in BMDM by Western blotting. β-actin levels were used for control of protein loading. The level of (F) IL-6, (G) TNF and (K) IL-10 levels were measured by ELISA assay in supernatant of BMDM cultures after 24h of stimulus. * *P* < 0.05 in comparison to the respective non-infected groups (MOCK) and # *P* < 0.05 versus respective Wt infected/stimulated group. Graphs are representative of three independent experiments.

Next, pharmacological data were further validated with macrophages from TLR4 knockout mice (TLR4-/-), and compared to macrophages from wild-type (WT) and TLR2 knockout (TLR2-/-) mice. In the absence of TLR4 signaling, neither IAV nor HA could induce necroptosis above the basal levels, as judged by the similar levels of LDH and annexinV^+^/PI^+^ cell counts between TLR4^-/-^ and mock-infected control **(Figure 2C-D and Figure S5)**. Similarly, to WT macrophages, cells lacking TLR2 underwent necroptosis due to exposure to IAV or HA **(Figure 2C-D and Figure S5)**.

To gain insight into the role of TLR4 in IAV/HA-triggered necroptosis, we analyzed the expression of p-RIPK1, RIPK3, and MLKL in WT and TLR4^-/-^ macrophages. We observed that the absence of TLR4 impaired the expression of p-RIPK1, RIPK3, and MLKL **(Figure 2E).** Moreover, the lack of TLR4 signaling prevented the increase of pro-inflammatory cytokines IL-6 **(Figure 2F)** and TNF **(Figure 2G)** induced by IAV, HA, or LPS (Figure 2E and F). On the other hand, the IL-10 levels were enhanced in the TLR4^-/-^ macrophages stimulated with LPS or infected with IAV **(Figure 2H)**, suggesting a stronger control of IAV-induced cytokine imbalance *in vitro*.

### IAV or its HA trigger necroptosis and pro-inflammatory response is a TNF-dependent manner in macrophages

Since IAV/HA induced cell death with the engagement of TLR4 signaling, we next evaluated whether TNF-α, a downstream pro-inflammatory cytokine produced during these events, could be involved in the necroptotic loop. IAV-infected or HA-exposed macrophages were pretreated with an anti-TNF-α neutralizing antibody, which prevented viral-induced necroptosis by 40-50 %, as shown by a reduction in annexinV^+^/PI^+^ cell counts **(Figure 3A-B)** and LDH levels **(Figure 3C)**. The IAV/HA-enhanced intracellular levels of RIPK1 were abolished by anti-TNF-α antibodies **(Figure 3D-E)**. By blocking TNF signaling, we also prevented the IAV/HA-induced production of nitric oxide **(Figure 3F)** and IL-6 **(Figure 3G)**. Notably, treatment with anti-TNF associated with increased IL-10 levels even in virus stimulated macrophages **(Figure 3H)**. Together, our *in vitro* results demonstrate that necroptosis induced by IAV/HA is triggered by a loop of events involving TLR4, TNF-α, p-RIPK1, leading to necroptosis and the exacerbation of inflammation **(Figure 3I)**.

**Figure 3.**
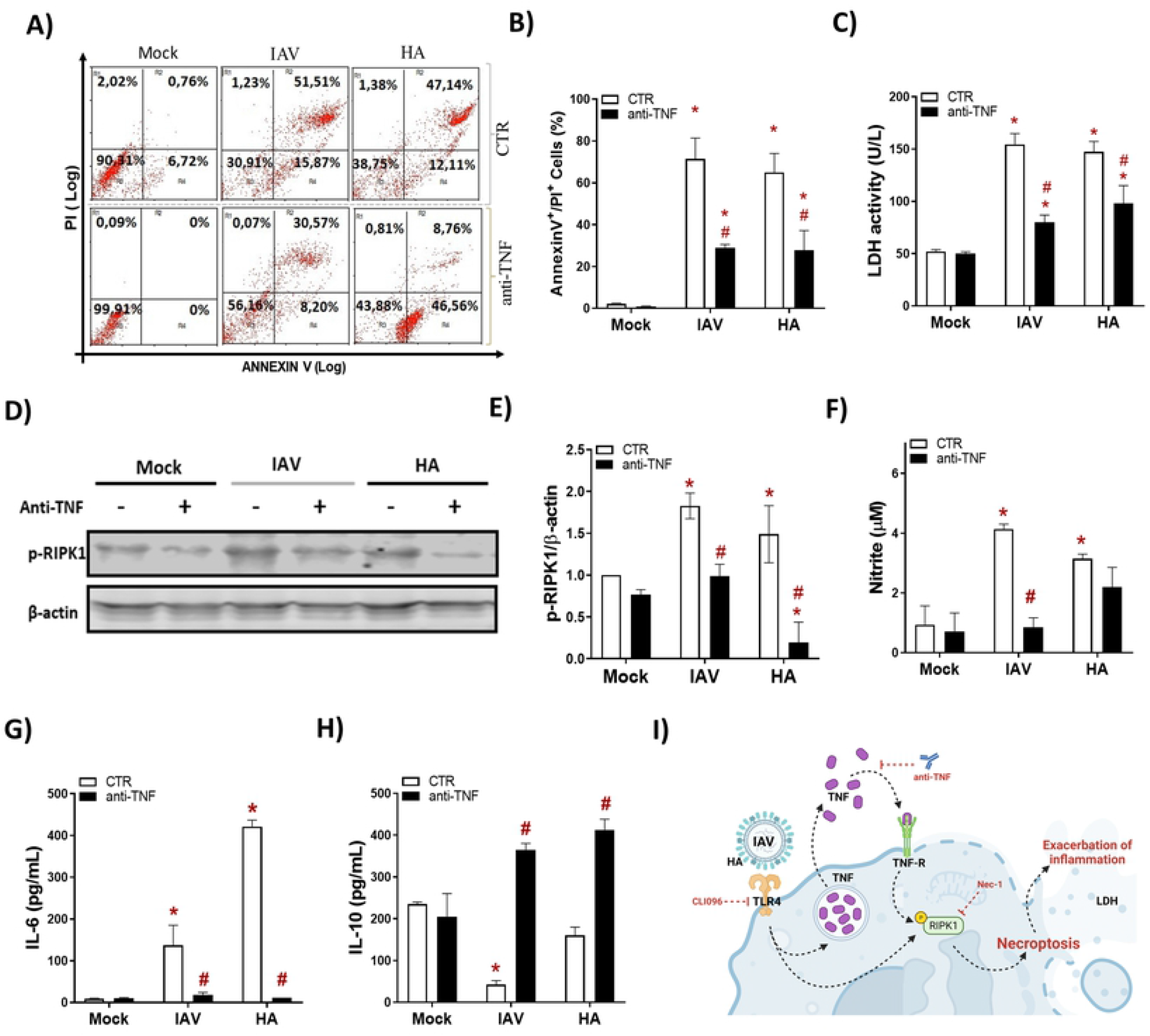
IAV- and HA-induced necroptosis is dependent a TNF. Macrophage cultures were pre-treated with 1 ng/mL anti-TNF-α antibody and then infected with IAV (MOI of 0.25) or exposed to 10 ng/mL of viral HA for 24 h. Cell death analysis was performed by flow cytometry analysis of annexinV/PI positive cells (A-B). Assessment of cell viability through the measurement of LDH release in the supernatant of BMDM (C). (D) The expression of p-RIPK1 were detected in BMDM by Western blotting. β-actin levels were used for control of protein loading. (E) Graphs of bands densitometry obtained after loading normalization and expressed as fold change over Mock untreated control. (F) Levels of nitrite were measured by Griess method in supernatant of BMDM cultures after 24h of stimulus. The levels of (H) IL-6 and (K) IL-10 were measured by ELISA assay in supernatant of BMDM cultures after 24h of stimulus. Data are presented as the mean ± SEM of 5 independent experiments * P < 0.05 versus untreated control group (MOCK); # P < 0.05 versus respective untreated infected/stimulated group.. (I) Model of A/HA-triggered necroptosis. IAV/HA-triggered necroptosis by a loop of events involving TLR4, TNF-α, RIPK1, leading to exacerbation of inflammation. Image created with Biorender.com.

### Blocking of IAV-induced necroptosis loop ameliorates severely infected animals

Our next step was to assess whether clinically approved Etanercept/ENBREL (anti-TNF-α biodrug) could impair the IAV-induced necroptotic loop. During the *in vivo* experiment, we observed that the IAV led to significant weight loss on the 3rd day post-infection (DPI) **(Figure 4A)**. In contrast, the Etanercept treatment at reference dose (2.5 mg/kg) resulted in the weight stabilization of IAV-infected mice, showing a significant difference compared to untreated ones from the 6 DPI afterwards **(Figure 4A)**. The protective effects of Etanercept were also observed in survival, because IAV-infected etanercept-treated mice presented significantly enhanced survival (by 8-times) when compared to untreated infected mice **(Figure 4B)**.

**Figure 4.**
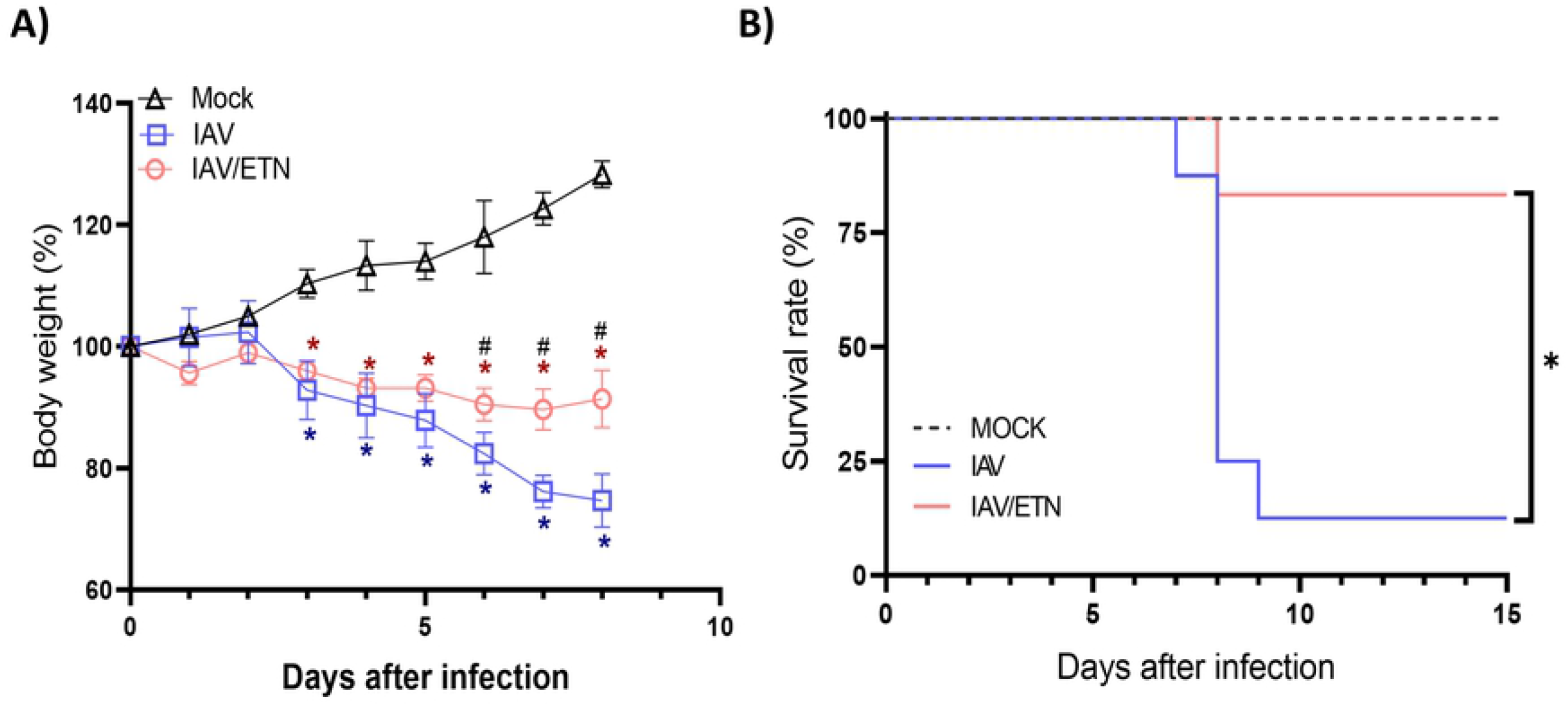
Etanercept reduced weigh loss improved survival and during IAV lethal infection in mice. C57Bl/6 mice were inoculated intranasally with 10^3^ PFU of IAV. Six hours post-infection, animals were treated intraperitoneally with of 2.5 mg/kg of Etanercept (ETN). Mice received a daily dose of ETN for 7 days and were monitored for survival **(A)** and body-weight loss **(B)** analysis. Data are show as percentage of survival and weight. Graphs are representative of three independent experiments. * *P* < 0.05 in comparison to IAV-infected untreated group.

We next analyzed the respiratory tract of the mice at 3 DPI and 5 DPI, before the mortality peak. Despite not inhibiting the total leukocyte accumulation in BAL **(Figure 5A-B)**, etanercept treatment significantly reduced the number of polymorphonuclear leukocytes (PMN) in both time points analyzed **(Figure 5A-B)**. Moreover, the etanercept-treated group presented a significant increase in the mononuclear leukocytes on 3 DPI. We observed that etanercept reduced the content of annexinV^+^/PI^+^ mononuclear cells induced by IAV, suggesting a necroptosis reduction during both times analyzed **(Figure 5C-D)**. Moreover, the etanercept-treated group also presented a reduction of IAV-induced overall cell death levels, as measured by LDH levels in BAL **(Figure 5E)**. In this context, we also evaluated whether the impairment of the TNF pathway could also reduce necroptosis in lung tissue. We observed an increased expression of pRIPK1 and MLKL in lung tissue of IAV-infected mice (**Figure 5F**). Treatment with etanercept prevented the increase of these protein levels **(Figure 5G-H)** while increased caspase-8 content, suggesting that anti-TNF treatment shift cell death from necroptosis to apoptosis.

**Figure 5:**
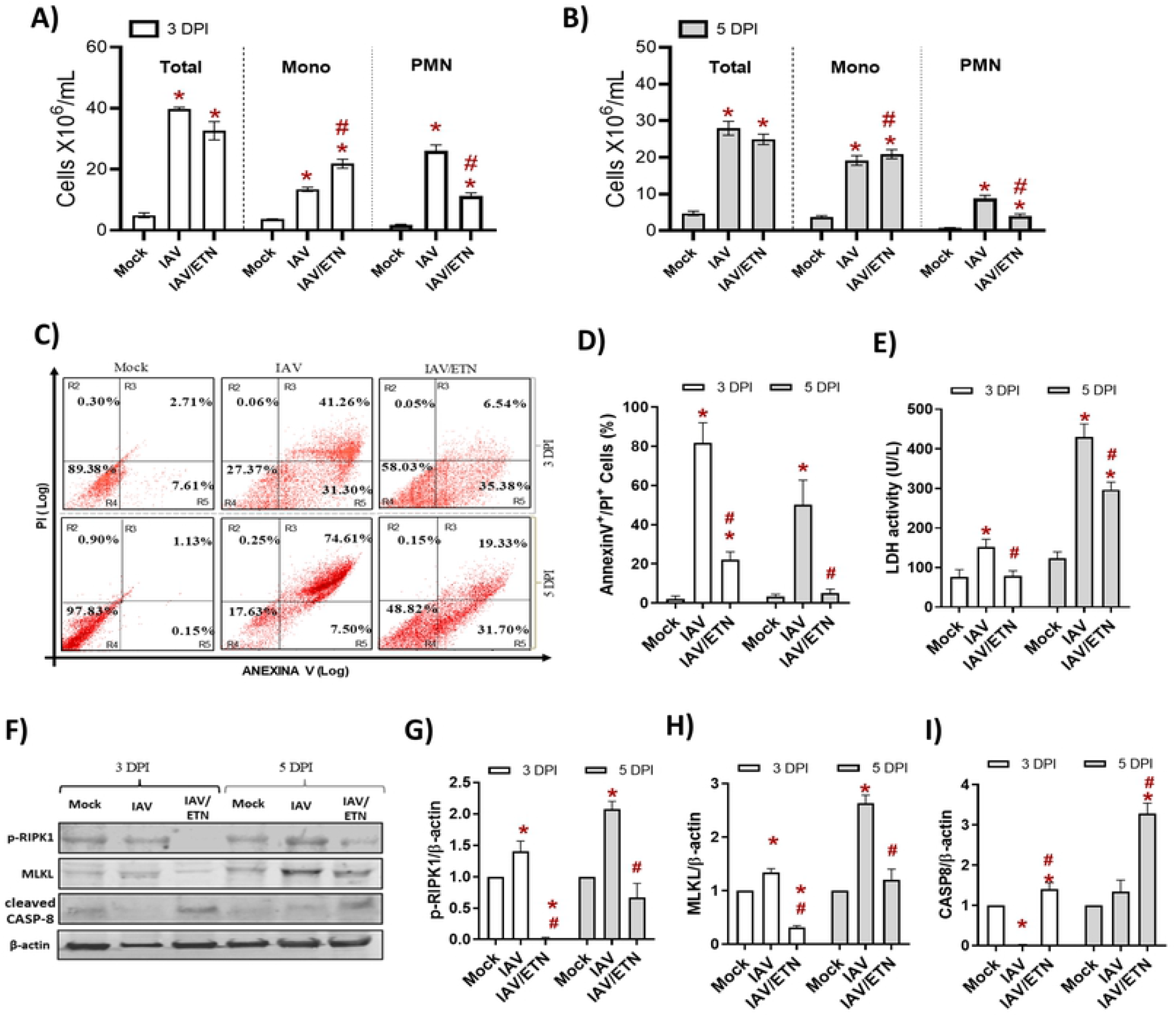
Etanercept reduced necroptosis *in vivo*. C57Bl/6 mice were inoculated intranasally with 10^3^ PFU of IAV with and without daily treatment with Etanercep (ETN, 2.5 mg/kg, i.p.). At days 3 and 5 post-infection, animals were euthanized, and BAL and lung were collected. **(A-B)** Total and differential cell counts in bronchoalveolar lavage (BAL) were represented as number of differential cell counts at days 3 **(A)** and 5 **(B)** post-infection. Total: leukocytes total, Mono: mononuclear leukocytes, PMN: polymorphonuclear leukocytes. **(C-D)** Mononuclear cells death was evaluated by annexinV/PI labelling. **(E)** Lung damage was evaluated by measuring LDH release in the bronchoalveolar lavage (BAL) **(F-I)** The expression of p-RIPK1, MLKL, and cleaved Caspase-8 were detected in the lung tissue homogenates by Western blotting. β-actin levels were used for control of protein loading). **(G-I)** Graphs of bands densitometry obtained after loading normalization and expressed as fold change over Mock control. Data are expressed as means ± SEM; One-way ANOVA with Dunnett’s post-hoc test. * *P* < 0.05 in comparison to Mock and # *P* < 0.05 in comparison to IAV-infected untreated group. Experiments were performed with 4-6 mice/group.

Treatment with etanercept reduced the levels lung inflammation in IAV-infected mice, as evaluated by protein content **(Figure 6A),** CXCL1/KC **(Figure 6B)**, CCL2/MCP1 **(Figure 6C)** IL-6 **(Figure 6D)**, TNF levels **(Figure 6E)**, IFN-γ **(Figure 6F)** and IL-10 **(Figure 6G)** levels in the BAL. Together, our data suggest that the impairment of a necroptosis loop ameliorates cytokines storms induced by IAV infection.

**Figure 6.**
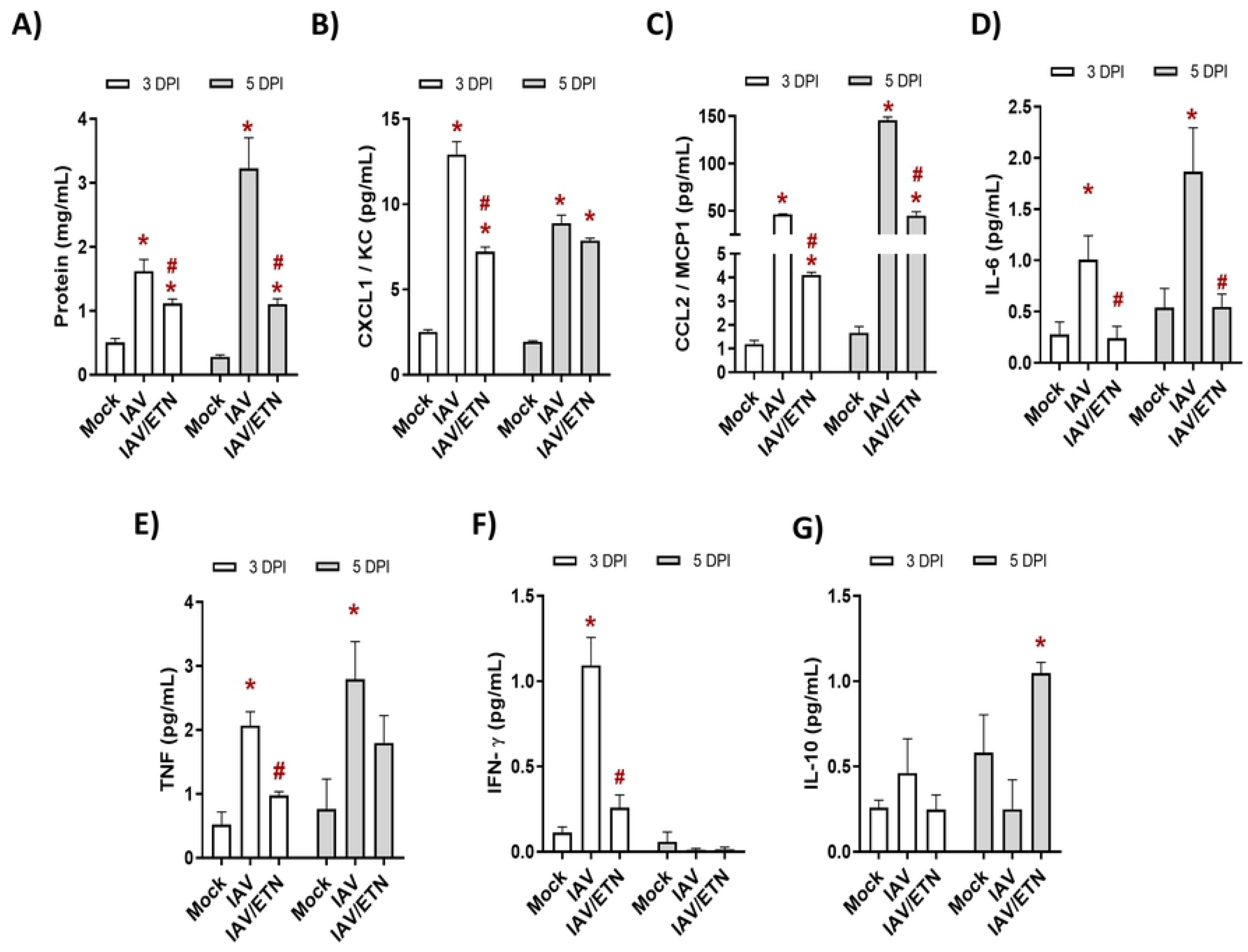
Etanercept reduced *cytokine storm* in IAV-infected mice. C57Bl/6 mice were inoculated intranasally with 10^3^ PFU of IAV with and without daily treatment with Etanercep (ETN, 2.5 mg/kg, i.p.). At days 3 and 5 post-infection (DPI), animals were euthanized and the bronchoalveolar lavage (BAL) was collected for quantification of the of total protein (A) and inflammatory mediators. The level of CXCL1/KC (B), CCL2/MCP1 (C), IL-6 (D), TNF (E), INF-γ (F) and IL-10 (G) were measured by ELISA. Data are expressed as means ± SEM; One-way ANOVA with Dunnett’s post-hoc test. * *P* < 0.05 in comparison to Mock and # *P* < 0.05 in comparison to IAV-infected untreated group. Experiments were performed with 4-6 mice/group.

## DISCUSSION

Emerging and re-emerging IAV strains are responsible for seasonal epidemics and pandemics, leading influenza to be a continuous warning to global public health systems [4,5]. One of the biggest challenges in the development of influenza antivirals is the difficulty to early provide treatments during acute viral infections to improve clinical outcomes [29]. Host-directed broad-spectrum antimicrobial drugs have been attempted to be used against respiratory viral infection, such as influenza and coronavirus disease 2019 (COVID-19) [30,31]. In this context, the use of immunomodulatory drugs has been reported as a promising approach for treating hypercytokinemia induced by acute viral infections [32].

After viral infection, an innate response is triggered in host cells to produce interferons, cytokines and chemokines to lead an immune activation, and in certain circumstances, the final fate of the host cell is its death [17]. The various ways by which cells die may directly influence viral physiopathology and patient’s clinical outcome [9,10]. With respect to pulmonary viral infections, it has been reported that different viruses lead to the induction of specific types of death mechanisms. Recently, we reported that pyroptosis is triggered by SARS-CoV-2 human monocytes, either by experimental infection and in critically ill COVID-19 patients [33]. For IAV, necroptosis and apoptosis have been reported in epithelial airways cells [11–14], where virus actively replicates. Here, we further described thatmacrophages, non-permissive cells to IAV replication, exposed to this virus or just its surface glycoprotein HA experience necroptosis, which leads to the imbalance of pro-inflammatory and regulatory modulators associated with cytokine storm and severe influenza *in vitro* and *in vivo*.

Indeed, necroptosis has been described as a highly inflammatory process [26], leading to lung dysfunction and the development of severe multiorgan tissue damage during virus infections [34–36]. Importantly, the inhibition of necroptosis in IAV/HA-exposed cells shifted macrophage from pro-inflammatory to regulatory phenotype [37], because IL-6 and nitric oxide levels were reduced and while IL-10 levels were enhanced.

We observed that TLR4, which plays a major role in the recognition of PAMPs [38–40], is engaged by IAV and its HA. Following this event, RIPK1 activation, RIPK3 engagement and MLKL expression are hallmarks of the pore-forming event that culminate with cell lysis^31^ measurable by quantification of PI+ cells and increased LDH levels detected throughout this study. Necroptosis is a TNF-dependent event [38,41] and, in fact, this pro-inflammatory cytokine was triggered during macrophage necroptosis provoked by IAV/HA. Moreover, the blockage of TNF by etanercept led to reduction of the expression of both p-RPK1 and MLKL in mononuclear cells *ex vivo*, while increasing the levels of cleaved caspase 8. Because caspase 8 is related to apoptosis, our data suggest that the *in vivo* intervention with etanercept on IAV-infected mice downregulated the necroptotic loop by altering the way monocular cellsdie and allowing the shift from necroptosis to apoptosis [38,42,43]

In summary, we demonstrated a positive feedback loop of events that led to necroptosis and exacerbated inflammation in IAV-infected macrophages. Our data demonstrated that IAV infection and its HA induce necrotic cell death by engaging TLR4 signaling to generate an enhanced TNF-α level and RIP1K/MLKL activation. Etanercept, which is endowed with anti-TNF-α activity, controlled severe influenza, impairing both necroptosis in IAV-infected mice. The etanercept-treated animals change their proinflammatory to a regulatory response, increasing survival. The present work improves the knowledge of influenza pathophysiology by highlighting the importance of macrophage cell death during severe infection.

## MATERIALS AND METHODS

### Reagents

We used different pharmacological inhibitors throughout this study (Table 1). The concentration of each inhibitor was chosen based on the manufacturer’s recommendations. All inhibitors were dissolved in 100% dimethylsulfoxide (DMSO) and subsequently diluted at least 10^4^-fold in culture medium before each assay. The final DMSO concentrations showed no cytotoxicity.

**Table 1.** List of small molecules and bio-drugs used as pharmacological inhibitors.

| Compound | Abbreviation | Target | Concentration | Manufacture | Catalog number |
| --- | --- | --- | --- | --- | --- |
| Z-VAD-FMK | ZVAD | Pan-caspase | 10 µM | Invivogen | tlrl-vad |
| Necrostatin-1 | Nec-1 | RIPK1 | 25 µM | Sigma-Andrich | 64419-92-7 |
| Recombinant Mouse TNF-alpha | TNF | TNF-receptor | 1 ng/mL | R&D System | 410-MT/CF |
| Resatorvid | CLI95 | TLR4 | 2 µM | Invivogen | tlrl-cli95 |
| anti-TNF antibody | anti-TNF | TNF | 1 ng/mL | R&D System | MAB4101 |
| Polymyxin b | POLI | LPS | 100 ng/mL | Sigma-Aldrich | P4932 |
| Lipopolysaccharide serotype 0111: B4 | LPS | TLR4 | 10ng/mL | Sigma-Aldrich | L4931 |
| Etanercept | ETN | TNF-receptor | 2.5 mg/kg | Pfizer |  |

Besides the pharmacological inhibitors, Recombinant Influenza A Virus Hemagglutinin H1 protein (HA, Abcam, Cat.# ab217651) was used. To select HA dose, macrophages were exposed to different concentrations of this protein or lipopolysaccharide (LPS; Sigma Aldrich). We observed that 10 ng/mL of HA was the minimal dose to induce TNF (Figure S1) and 10 ng/mL of LPS was used as positive control **(Figure S1)**.

### Virus Strain and Growth Conditions

Madin-Darby Canine Kidney (MDCK) were cultured in Dulbecco’s Modified Eagle Medium (DMEM; Life Technologies) and supplemented with 10 % fetal bovine serum (FBS; HyClone), 100 U/mL penicillin, and 100 mg/mL streptomycin (Sigma-Aldrich) at 37 °C in a 5 % CO_2_ atmosphere.

IAV, A/H1N1/Puerto Rico/8/34 strain (PR8), was grown and titrated in MDCK [44,45]. Briefly, confluent cells in 75 cm^2^ culture flasks were infected with PR8 at the multiplicity of infection (MOI) of 0.1. The inoculum was prepared in DMEM containing 2 μg/ml of TPCK-trypsin (Sigma-Aldrich). Cells were exposed to PR8 for 1 h at 37 °C to allow virus adsorption. After the incubation period, cells were washed with PBS, and fresh inoculation medium was added. Cultures were accompanied daily until the detection of IAV-induced cytopathic effect (CPE). Culture supernatants containing the viruses were collected and centrifuged (1,500 rpm for 5 minutes) to remove cell debris. The viral stocks were aliquoted and stored at −70 °C for further studies.

Titration was performed through plaque assays, using 10-fold dilutions (10^-1^ to 10^-10^) in 6-well plates. After 1h incubation with virus dilutions, the inoculum was removed, cells were washed, and fresh DMEM with 2.5% agarose with TPCK-trypsin at 1 μg/ml was added. After 3 days at 37 °C, laying agar was removed, the wells were washed with PBS and stained with crystal violet 0.1 %.

### Mice

C57BL/6 mice (20–30 g) were supplied by the Institute of Science and Technology in Biomodels from Oswaldo Cruz Foundation and used at 8–12 weeks of age. The Institutional animal welfare committee (Committee on the Use of Laboratory Animals of the Oswaldo Cruz Foundation) approved all animal experiments in agreement with the Brazilian National guidelines supported by CONCEA (Conselho Nacional de Controle em Experimentação Animal) under license number L050/15 (CEUA/FIOCRUZ). Mice were maintained with rodent diet and water available *ad libitum* with 12h light-dark cycle under controlled temperature (23 ± 1 °C).

### Murine and human macrophages

To obtain murine bone marrow-derived macrophages (BMDM), cells isolated from femur and tibia of mice with C57BL/6 background; either from wild type (WT), Toll-like Receptor 2 or 4 knockout (TLR2^-/-^ or TLR4^-/-^, respectively) mice. The isolated cells were cultured for 7 days in RPMI-1640 medium supplemented with 30% (vol/vol) L929 supernatant, 20% (vol/vol) heat-inactivated fetal bovine serum, 1% L-glutamine (vol/vol), and 1% penicillin-streptomycin (vol/vol) as previously described by Assunção et al., (2017). Differentiated macrophages were cultured in RPMI-1640 supplemented with 10% heat-inactivated fetal bovine serum (vol/vol), 1% L-glutamine (vol/vol), and 1% penicillin/streptomycin (vol/vol). BMDM cells were maintained at a density of 5×10^5^ cells/ml.

Human monocyte-derived macrophages (MDMs) were obtained through plastic adherence of peripheral blood mononuclear cells (PBMCs). In brief, PBMCs were obtained from buffy coat preparations of healthy donors by density gradient centrifugation (Ficoll-Paque, GE Healthcare). PBMCs (2.0 x 10^6^ cells) were plated onto 48-well plates (NalgeNunc) in DMEM containing 10 % human serum (HS; Millipore) and penicillin/streptomycin. Cells were maintained for monocyte differentiation into macrophages at standard culture conditions for 6–7 days. Then, non-adherent cells were washed, and the remaining macrophage layer was maintained in DMEM with 5 % HS [47].

The purity of murine and human macrophages was above 95 %, as determined by flow cytometric analysis (FACScan; Becton Dickinson) using anti-CD3 (BD Biosciences) and anti-CD16 (Southern Biotech) monoclonal antibodies.

### In vitro experiments

Murine or human macrophages were plated (5 x 10^5^ cells/well) in 24-well culture plates (flat-bottom, tissue-culture-treated plates; Costar) and were incubated for 12 h at 37°C and 5% CO_2_. The cultures were then infected with PR8 (MOI of 0.25) or exposed to 10 ng/mL of HA for 24 h. In parallel, as positive controls, macrophage cultures were also stimulated with 1 ng/mL of TNF-α or 10 ng/mL of LPS. After this incubation period, culture supernatants were collected for cell death analysis by LDH measurement and cytokines/chemokines quantification. The cell monolayers were harvested for flow cytometry and western blot analysis. To impair the cell death, 10 μM of zVAD, 25 μM of Nec-1, 2 μM of CLI95, or 1 ng/mL of anti-TNF-α antibody were added to the cell culture and incubated for 30 min before IAV-infection or the application of stimuli (HA, LPS or TNF), and remained for all the infection/stimulus time, at 37°C in 5% CO_2_.

### Analysis of cell death

Macrophage’s viability was evaluated by quantifying LDH and by flow cytometry in the presence of the pharmacological inhibitors. LDH quantification was performed after culture supernatants centrifugation (5,000 rpm for 1 minute) to remove cellular debris. According to the manufacturer’s instructions, extracellular lactate dehydrogenase (LDH) was quantified using the Doles^®^ kit. In summary, 25 μL of cell samples were seeded in 96-well culture plates and incubated with 5 μL of ferric alum and 100 μL of LDH substrate for 3 minutes at 37 °C. Nicotinamide adenine dinucleotide (NAD, oxidized form) was added, followed by a stabilizing solution. After 10 minutes-incubation, plates were read in a spectrophotometer at 492 nm.

For flow cytometry analysis, macrophages were diluted in a labeling buffer (10^6^ cells/mL). Then, 100 μL of cell samples were marked with 5 μL of AnnexinV (BD Biosciences) and 1 μL of PI (BD Biosciences) for 15 minutes for cell death analysis. Around 10,000 events were acquired using FACSCalibur, and analyses were performed using the CellQuest software. Macrophages were gated through cell size (forward light scatter, FSC) and granularity (side light scatter, SSC) analysis (Figure S2). The profiles for macrophage’s positivity to AnnexinV and/or PI (AnnexinV^+^/PI^+^) were determined for cells from *in vitro* experiments and from BAL of IAV-infected mice. Data acquisition was set to count a total of 10,000 events, and the FLOWJO software package was used to analyze the data.

### In vivo experiments

For infection procedures, mice were anesthetized with 60 mg/kg of ketamine and 4 mg/kg of xylazine and inoculated intranasally with PBS (MOCK) or 10^3^ PFU of PR8 in 25 μl of PBS [48]. The animals were kept under observation until they completely recovered. Six hours post-infection, the treated groups received an intraperitoneal dose of 2.5 mg/kg of Etanercept/ENBREL in 200 μL of vehicle (PBS). The treatment was continued with a daily dose of 2,5 mg/kg of etanercept for 7 days. We used 10 mice per experimental group: mock-infected (MOCK); influenza-infected and treated with vehicle (IAV); and influenza-infected and treated with Etanercept (IAV/ETN). The animals were monitored daily for 15 days for survival and eight days for body-weight analysis. In the case of weight loss higher than 25 %, euthanasia was performed to alleviate animal suffering.

### Bronchoalveolar lavage and lung homogenates

Mice were euthanized on days 3 and 5 after infection to evaluate the lung’s inflammatory process induced by IAV infection. The mice were anesthetized, and bronchoalveolar lavage (BAL) from both lungs was harvested by washing the lungs three times with two 1-ml aliquots of cold PBS. After centrifugation of BAL (1500 rpm for 5 minutes), the pellet was used for total and differential leukocytes counts and cell death analysis by flow cytometry. The supernatant of the centrifuged BAL was used for cytokines/chemokines and total protein measurements and cell death analysis by LDH quantification. Total leukocytes (diluted in Turk’s 2 % acetic acid fluid) were counted using a Neubauer chamber. Differential cell counts were performed in cytospins (Cytospin3; centrifugation of 350 x *g* for 5 minutes at room temperature) and stained by the May-Grünwald-Giemsa method. The levels of cytokines and chemokines were assessed by ELISA. The total protein concentration in the BAL was measured using a BCA protein assay kit (Thermo Scientific).

After BAL harvesting, the lungs were perfused with 5 ml of PBS to remove the circulating blood. Lungs were then collected and macerated in 750 μL of cold phosphate buffer containing protease inhibitor cocktail (Roche Applied Science, Mannheim, Germany). Homogenates were stored at −80 °C for western blot analysis.

### Measurements Inflammatory Mediators

The levels of TNF, IL-6, IL-10, IFN-γ, CCL2 or CXCL1 were quantified in the *in vitro* macrophage supernatants and BAL from IAV-infected mice using DuoSet® ELISA assays, following the manufacturer’s instructions (R&D Systems). Briefly, 100 μL of each sample were added in 96-well plates covered with the capture antibody. After a 2 h-incubation period at room temperature (RT), the detection antibody was added, and the plates were incubated for a second round of 2 h at RT. Streptavidin-HRP and its substrate were added with a 20 min incubation interval, and the optical density was determined using a microplate reader set to 450 nm. Nitrite levels in cell-free culture supernatant were measured using the Griess reagent system according to the manufacturer’s instructions (Promega cat.# G2930).

### Western blot assay

Cellular extracts of 1×10^6^ cells or 0.5 g of tissue were homogenized in the RIPA lysis buffer (1% Triton X-100, 2% SDS, 150 mM NaCl, 10 mM HEPES, 2 mM EDTA) containing protease and phosphatase inhibitor cocktail (Roche, pH 8.0). After centrifugation at 13 000 g for 5 min, cell lysates were prepared in reducing and denaturing conditions and subjected to SDS-PAGE. Equal concentrations of proteins were fractionated by electrophoresis on 10% of acrylamide gels. The proteins were transferred onto a nitrocellulose membrane (Millipore, Billerica, MA, USA), followed by blocking of nonspecific binding sites in 5% nonfat milk in TBST (50 mM Tris-HCl - pH 7.4, 150 mM NaCl, 0.05% Tween 20) for 1 h at room temperature and blotted with primary antibodies in TBST overnight at 4 °C. The following antibodies were used: anti-Phospho-RIPK1 (Ser166) (Cell Signaling- # 31122S), anti-RIPK3 (D8J3L) (Cell Signaling - # 15828S), anti-MLKL (D2I6N) (Cell Signaling- #14993S), and anti-β-actin (Sigma, #A1978) Proteins of interest were identified by incubating the membrane with IRDye® LICOR secondary antibodies in TBST, followed by fluorescence imaging detection using the Odyssey® system (CLx Imaging System). Protein bands were quantified by densitometric image analysis using the ImageJ software. All the data were normalized by β-actin expression quantification.

### Statistical analysis

All experiments were carried out at least three times independently, including technical replicates in each assay. Statistical analysis was carried out using the GraphPad Prism software. *P* values were calculated by unpaired Student’s t test, except for PMC calculated with Wilcoxon rank-sum test. Results are expressed as mean ± SEM (median (IQR)). The significance of the survival curves was evaluated using the Log-rank (Mantel-Cox) test. *P* values < 0.05 were considered statistically significant.

## Acknowledgements

We would like to thank Dra. Juliana De Meis from Laboratório de Pesquisa sobre o Timo/FIOCRUZ for kindly donated etanercept (ENBREL®) and Dr. Marcelo Bozza from Instituto de Microbiologia Paulo de Goes/Federal University of Rio de Janeiro, for donating C57Bl/6 knockout mice. We thank to Fiocruz Luminex Platform (*Subunidade Luminex-RPT03C* Rede de Plataformas PDTIS, FIOCRUZ/RJ) and the assistance of MSc Edson Fernandes de Assis for the use of its Luminex facilities.

## References

1. Webster RG, Bean WJ, Gorman OT, Chambers TM, Kawaoka Y. Evolution and ecology of influenza A viruses. Microbiol Rev. 1992;56: 152–179. doi:10.1128/mr.56.1.152-179.1992

2. Fineberg H V. Pandemic Preparedness and Response — Lessons from the H1N1 Influenza of 2009. N Engl J Med. 2014;370: 1335–1342. doi:10.1056/NEJMRA1208802

3. Paules C, Subbarao K. Influenza. Lancet. 2017;390: 697–708. doi:10.1016/S0140-6736(17)30129-0

4. World Health Organization. Global influenza strategy 2019-2030. World Health Organization, editor. World Health Organization. Geneva: World Health Organization; 2019. Available: https://apps.who.int/iris/handle/10665/311184.

5. Iuliano AD, Roguski KM, Chang HH, Muscatello DJ, Palekar R, Tempia S, et al. Estimates of global seasonal influenza-associated respiratory mortality: a modelling study. Lancet. 2018;391: 1285–1300. doi:10.1016/S0140-6736(17)33293-2

6. Paget J, Spreeuwenberg P, Charu V, Taylor RJ, Iuliano AD, Bresee J, et al. Global mortality associated with seasonal influenza epidemics: New burden estimates and predictors from the GLaMOR Project. J Glob Health. 2019;9: 1–12. doi:10.7189/jogh.09.020421

7. Resa-Infante P, Jorba N, Coloma R, Ortin J. The influenza virus RNA synthesis machine. RNA Biol. 2011;8: 207–215. doi:10.4161/rna.8.2.14513

8. Kalil AC, Thomas PG. Influenza Fisiopatología. Crit Care. 2019;23: 1–7.

9. Yatim N, Albert ML. Dying to Replicate: The Orchestration of the Viral Life Cycle, Cell Death Pathways, and Immunity. Immunity. 2011;35: 478–490. doi:10.1016/j.immuni.2011.10.010

10. Nailwal H, Chan FK-M. Necroptosis in anti-viral inflammation. Cell Death Differ. 2019;26: 4–13. doi:10.1038/s41418-018-0172-x

11. Kuriakose T, Man SM, Subbarao Malireddi RK, Karki R, Kesavardhana S, Place DE, et al. ZBP1/DAI is an innate sensor of influenza virus triggering the NLRP3 inflammasome and programmed cell death pathways. Sci Immunol. 2016;1. doi:10.1126/sciimmunol.aag2045

12. Nogusa S, Thapa RJ, Dillon CP, Liedmann S, Oguin TH, Ingram JP, et al. RIPK3 Activates Parallel Pathways of MLKL-Driven Necroptosis and FADD-Mediated Apoptosis to Protect against Influenza A Virus. Cell Host Microbe. 2016;20: 13–24. doi:10.1016/j.chom.2016.05.011

13. Thapa RJ, Ingram JP, Ragan KB, Nogusa S, Boyd DF, Benitez AA, et al. DAI Senses Influenza A Virus Genomic RNA and Activates RIPK3-Dependent Cell Death. Cell Host Microbe. 2016;20: 674–681. doi:10.1016/j.chom.2016.09.014

14. Zhang T, Yin C, Boyd DF, Quarato G, Ingram JP, Shubina M, et al. Influenza Virus Z-RNAs Induce ZBP1-Mediated Necroptosis. Cell. 2020;180: 1115–1129.e13. doi:10.1016/j.cell.2020.02.050

15. Atkin-Smith GK, Duan M, Chen W, Poon IKH. The induction and consequences of Influenza A virus-induced cell death. Cell Death Dis. 2018;9. doi:10.1038/s41419-018-1035-6

16. Shubina M, Tummers B, Boyd DF, Zhang T, Yin C, Gautam A, et al. Necroptosis restricts influenza A virus as a stand-alone cell death mechanism. J Exp Med. 2020;217. doi:10.1084/jem.20191259

17. Orzalli MH, Kagan JC. Apoptosis and Necroptosis as Host Defense Strategies to Prevent Viral Infection. Trends Cell Biol. 2017;27: 800–809. doi:10.1016/j.tcb.2017.05.007

18. Yang Y, Tang H. Aberrant coagulation causes a hyper-inflammatory response in severe influenza pneumonia. Cell Mol Immunol. 2016;13: 432–442. doi:10.1038/cmi.2016.1

19. Marvin SA, Russier M, Huerta CT, Russell CJ, Schultz-Cherry S. Influenza Virus Overcomes Cellular Blocks To Productively Replicate, Impacting Macrophage Function. Dermody TS, editor. J Virol. 2017;91. doi:10.1128/JVI.01417-16

20. Sakabe S, Iwatsuki-Horimoto K, Takano R, Nidom CA, Le M thi Q, Nagamura-Inoue T, et al. Cytokine production by primary human macrophages infected with highly pathogenic H5N1 or pandemic H1N1 2009 influenza viruses. J Gen Virol. 2011;92: 1428–1434. doi:10.1099/vir.0.030346-0

21. Tisoncik JR, Korth MJ, Simmons CP, Farrar J, Martin TR, Katze MG. Into the Eye of the Cytokine Storm. Microbiol Mol Biol Rev. 2012;76: 16–32. doi:10.1128/MMBR.05015-11

22. Meischel T, Villalon-Letelier F, Saunders PM, Reading PC, Londrigan SL. Influenza A virus interactions with macrophages: Lessons from epithelial cells. Cell Microbiol. 2020;22: 1–11. doi:10.1111/cmi.13170

23. Frabutt DA, Wang B, Riaz S, Schwartz RC, Zheng Y-H. Innate Sensing of Influenza A Virus Hemagglutinin Glycoproteins by the Host Endoplasmic Reticulum (ER) Stress Pathway Triggers a Potent Antiviral Response via ER-Associated Protein Degradation. Schultz-Cherry S, editor. J Virol. 2018;92. doi:10.1128/JVI.01690-17

24. Liu H, Ma Y, Pagliari LJ, Perlman H, Yu C, Lin A, et al. TNF-α-Induced Apoptosis of Macrophages Following Inhibition of NF-κB: A Central Role for Disruption of Mitochondria. J Immunol. 2004;172: 1907–1915. doi:10.4049/jimmunol.172.3.1907

25. Wu YT, Tan HL, Huang Q, Sun XJ, Zhu X, Shen HM. ZVAD-induced necroptosis in L929 cells depends on autocrine production of TNFα mediated by the PKC-MAPKs-AP-1 pathway. Cell Death Differ. 2011;18: 26–37. doi:10.1038/cdd.2010.72

26. Dhuriya YK, Sharma D. Necroptosis: A regulated inflammatory mode of cell death. J Neuroinflammation. 2018;15: 1–9. doi:10.1186/s12974-018-1235-0

27. Liu W-C, Lin S-C, Yu Y-L, Chu C-L, Wu S-C. Dendritic Cell Activation by Recombinant Hemagglutinin Proteins of H1N1 and H5N1 Influenza A Viruses. J Virol. 2010;84: 12011–12017. doi:10.1128/jvi.01316-10

28. Zhang S, Gu D, Ouyang X, Xie W. Proinflammatory effects of the hemagglutinin protein of the avian influenza A (H7N9) virus and microRNA-mediated homeostasis response in THP-1 cells. Mol Med Rep. 2015;12: 6241–6246. doi:10.3892/mmr.2015.4142

29. Muthuri SG, Venkatesan S, Myles PR, Leonardi-Bee J, Al Khuwaitir TSA, Al Mamun A, et al. Effectiveness of neuraminidase inhibitors in reducing mortality in patients admitted to hospital with influenza A H1N1pdm09 virus infection: a meta-analysis of individual participant data. Lancet Respir Med. 2014;2: 395–404. doi:10.1016/S2213-2600(14)70041-4

30. Li CC, Wang XJ, Wang HCR. Repurposing host-based therapeutics to control coronavirus and influenza virus. Drug Discov Today. 2019;24: 726–736. doi:10.1016/j.drudis.2019.01.018

31. Pan H, Peto R, Henao-Restrepo A, Preziosi M, Sathi-yamoorthy V, Abdool Karim Q, et al. Repurposed Antiviral Drugs for Covid-19 — Interim WHO Solidarity Trial Results. N Engl J Med. 2021;384: 497–511. doi:10.1056/NEJMoa2023184

32. Gu Y, Zuo X, Zhang S, Ouyang Z, Jiang S, Wang F, et al. The Mechanism behind Influenza Virus Cytokine Storm. Viruses. 2021;13: 1362. doi:10.3390/v13071362

33. Ferreira AC, Soares VC, de Azevedo-Quintanilha IG, Dias S da SG, Fintelman-Rodrigues N, Sacramento CQ, et al. SARS-CoV-2 engages inflammasome and pyroptosis in human primary monocytes. Cell Death Discov. 2021;7: 43. doi:10.1038/s41420-021-00428-w

34. De Jong MD, Simmons CP, Thanh TT, Hien VM, Smith GJD, Chau TNB, et al. Fatal outcome of human influenza A (H5N1) is associated with high viral load and hypercytokinemia. Nat Med. 2006;12: 1203–1207. doi:10.1038/nm1477

35. McGonagle D, O’Donnell JS, Sharif K, Emery P, Bridgewood C. Immune mechanisms of pulmonary intravascular coagulopathy in COVID-19 pneumonia. Lancet Rheumatol. 2020;2: e437–e445. doi:10.1016/S2665-9913(20)30121-1

36. Fajgenbaum DC, June CH. Cytokine Storm. N Engl J Med. 2020;383: 2255–2273. doi:10.1056/nejmra2026131

37. O’Neill LAJ, Kishton RJ, Rathmell J. A guide to immunometabolism for immunologists. Nat Rev Immunol. 2016;16: 553–565. doi:10.1038/nri.2016.70

38. Bertheloot D, Latz E, Franklin BS. Necroptosis, pyroptosis and apoptosis: an intricate game of cell death. Cell Mol Immunol. 2021;18: 1106–1121. doi:10.1038/s41423-020-00630-3

39. He S, Liang Y, Shao F, Wang X. Toll-like receptors activate programmed necrosis in macrophages through a receptor-interacting kinase-3-mediated pathway. Proc Natl Acad Sci U S A. 2011;108: 20054–20059. doi:10.1073/pnas.1116302108

40. Kaiser WJ, Sridharan H, Huang C, Mandal P, Upton JW, Gough PJ, et al. Toll-like Receptor 3-mediated Necrosis via TRIF, RIP3, and MLKL. J Biol Chem. 2013;288: 31268–31279. doi:10.1074/jbc.M113.462341

41. Orzalli MH, Kagan JC. Apoptosis and Necroptosis as Host Defense Strategies to Prevent Viral Infection. Trends Cell Biol. 2017;27: 800–809. doi:10.1016/j.tcb.2017.05.007

42. Someda M, Kuroki S, Miyachi H, Tachibana M, Yonehara S. Caspase-8, receptor-interacting protein kinase 1 (RIPK1), and RIPK3 regulate retinoic acid-induced cell differentiation and necroptosis. Cell Death Differ. 2020;27: 1539–1553. doi:10.1038/s41418-019-0434-2

43. Newton K, Wickliffe KE, Dugger DL, Maltzman A, Roose-Girma M, Dohse M, et al. Cleavage of RIPK1 by caspase-8 is crucial for limiting apoptosis and necroptosis. Nature. 2019;574: 428–431. doi:10.1038/s41586-019-1548-x

44. Szretter KJ, Balish AL, Katz JM. Influenza: Propagation, Quantification, and Storage. Curr Protoc Microbiol. 2006;3: 1–22. doi:10.1002/0471729256.mc15g01s3

45. Organization. WH. Manual for the laboratory diagnosis and virological surveillance of influenza. Geneva; 2011. Available: https://apps.who.int/iris/handle/10665/44518

46. Assunção LS, Magalhães KG, Carneiro AB, Molinaro R, Almeida PE, Atella GC, et al. Schistosomal-derived lysophosphatidylcholine triggers M2 polarization of macrophages through PPARγ dependent mechanisms. Biochim Biophys Acta - Mol Cell Biol Lipids. 2017;1862: 246–254. doi:10.1016/j.bbalip.2016.11.006

47. Mesquita M, Fintelman-Rodrigues N, Sacramento CQ, Abrantes JL, Costa E, Temerozo JR, et al. HIV-1 and Its gp120 Inhibits the Influenza A(H1N1)pdm09 Life Cycle in an IFITM3-Dependent Fashion. Tompkins SM, editor. PLoS One. 2014;9: e101056. doi:10.1371/journal.pone.0101056

48. Barbosa R, Salgado A, Garcia CC, Filho BG, Apdf G. Protective Immunity and Safety of a Genetically Modified Influenza Virus Vaccine. PLoS One. 2014;9: 98685. doi:10.1371/journal.pone.0098685

